# RETINOID ORPHAN RECEPTOR GAMMA T (RORγT) PROMOTES INFLAMMATORY EOSINOPHILIA BUT IS DISPENSABLE FOR INNATE IMMUNE-MEDIATED COLITIS

**DOI:** 10.1101/2023.10.27.564391

**Authors:** Alvaro Torres-Huerta, Katelyn Ruley-Haase, Theodore Reed, Antonia Boger-May, Derek Rubadeux, Lauren Mayer, Arpitha Mysore Rajashekara, Morgan Hiller, Madeleine Frech, Connor Roncagli, Cameron Pederson, Mary Catherine Camacho, Lauren Hollmer, Lauren English, Grace Kane, David L. Boone

**Author notes:** These authors contributed equally to this work (Alvaro Torres-Huerta, Katelyn Ruley-Haase).

## Abstract

Inflammatory bowel diseases (IBD) result from uncontrolled inflammation in the intestinal mucosa leading to damage and loss of function. Both innate and adaptive immunity contribute to the inflammation of IBD and innate and adaptive immune cells reciprocally activate each other in a forward feedback loop. In order to better understand innate immune contributions to IBD, we developed a model of spontaneous 100% penetrant, early onset colitis that occurs in the absence of adaptive immunity by crossing villin-TNFAIP3 mice to RAG1^−/−^ mice (TRAG mice). This model is driven by microbes and features increased levels of innate lymphoid cells in the intestinal mucosa. To investigate the role of type 3 innate lymphoid cells (ILC3) in the innate colitis of TRAG mice, we crossed them to retinoid orphan receptor gamma t deficient (Rorγt^−/−^) mice. Rorγt^−/−^ x TRAG mice exhibited markedly reduced eosinophilia in the colonic mucosa, but colitis persisted in these mice. Colitis in Rorγt^−/−^ x TRAG mice was characterized by increased infiltration of the intestinal mucosa by neutrophils, inflammatory monocytes, macrophages and other innate cells. RNA and cellular profiles of Rorγt^−/−^ x TRAG mice were consistent with a lack of ILC3 and ILC3 derived cytokines, reduced antimicrobial factors, increased activation oof epithelial repair processes and reduced activation of epithelial cell STAT3. The colitis in Rorγt^−/−^ x TRAG mice was ameliorated by antibiotic treatment indicating that microbes contribute to the ILC3-independent colitis of these mice. Thus, Rorγt promotes eosinophilia but Rorγt and Rorγt-dependent ILC3 are dispensable for the innate colitis in TRAG mice.

## INTRODUCTION

Inflammatory bowel diseases (Crohn’s disease and ulcerative colitis) are characterized by damaging chronic inflammation in the gastrointestinal tract. The causes of inflammatory bowel diseases (IBD) are not known but a current paradigm is that genetics and environment contribute to susceptibility to disease[1, 2]. The environmental factors that contribute to IBD are not known but it is likely that these factors manifest, at least in part, through changes in the gut microbiome. A key component of host interactions with the microbial world is the innate immune system and so innate immunity is considered central to the pathogenesis of IBD[3].

Both innate and adaptive immunity contribute to IBD. Innate cells detect microbes and initiate the immune response, leading to activation and expansion of antigen specific adaptive immune cells. The adaptive responses produce cytokines that subsequently activate innate defenses, which can further induce adaptive immune cells in a positive feedback loop. This reciprocal interaction between innate and adaptive immunity creates the inflammation needed to defend against pathogens, but in pathological states like IBD the inflammation persists and damages the gut tissue leading to loss of function and IBD. A better understanding the causes of IBD requires more complete information about the nature of both the adaptive and innate immune pathology. However, the reciprocal nature of the adaptive and innate feedback loop makes it challenging to identify innate mechanisms that lead to IBD. Innate models of IBD that occur in the absence of adaptive immunity can address this challenge and provide insight into innate mechanisms of inflammation[4-6]. For example, a key role for type 3 innate lymphoid cells in IBD was revealed by studies using innate models of IBD lacking the transcription factor retinoid orphan receptor gamma t (RoRγt)[5]. Thus, models of IBD that occur in the absence adaptive immunity can provide valuable insight into innate immune mechanisms of disease.

We have developed a unique innate immune model of IBD that occurs spontaneously with 100% penetrance at 6 weeks-of-age in C57Bl/6 mice[6, 7]. Transgenic mice expressing tumor necrosis factor induced protein 3 (TNFAIP3; aka A20) in intestinal epithelial cells under the villin promoter (v-TNFAIP3) develop colitis on the recombination activating 1 (RAG1) deficient background (v-TNFAIP3 x RAG1^−/−^)[6]. The mucosal inflammation in v-TNFAIP3 x RAG1^−/−^ (TRAG) mice is evident as increased innate immune infiltration in the colon by 4 weeks of age and histologically by 6 to 8 weeks of age[6]. Antibiotics prevent colitis in TRAG mice, indicating that microbes drive the colitis in this innate model[6]. The expression of TNFAIP3 in the intestinal epithelium leads to altered microbial biogeography with invasion of the inner mucus layer by microbes, predominantly of the class Actinobacteria and Gammaproteobacteria[7, 8]. Despite this microbial invasion, v-TNFAIP3 mice do not develop colitis[7-9]. RAG1^−/−^ mice maintain an inner mucus layer largely devoid of microbes and also do not develop colitis[7]. However, 100% of TRAG mice develop colitis indicating that a combination of microbial invasion of the inner mucus layer and immunodeficiency leads to innate immune colitis. We found that treatment of TRAG mice with anti-Thy1.2 antibody to deplete innate lymphoid cells (ILC) could both prevent or reverse established colitis[6]. However, unlike other innate models of IBD, crossing TRAG mice to mice lacking RoRγt did not prevent colitis[6]. Thus, colitis in TRAG mice likely involves unique innate immune mechanisms and so herein we provide a more detailed characterization of the colitis that occurs in TRAG mice in the absence of RoRγt.

## MATERIALS AND METHODS

### Animal studies

Mice were bred and housed in the Freimann Life Sciences Center at the University of Notre Dame. All protocols were performed at the University of Notre Dame Freimann Life Science Center approved by Institutional Animal Care and Use Committees. All mice were bred and maintained on the C57BL/6 background. The villin-TNFAIP3 strain was generated previously using BAC-recombineering of the villin locus and characterized as described[9]. RAG1^−/−^ and Rorc^−/−^ mice (C57Bl/6) were purchased from Jackson Laboratories and interbred to villin-TNFAIP3 mice to generate villin-TNFAIP3 × RAG1^−/−^ (TRAG) mice, Rorc^−/−^ x TRAG mice, or RAG1^−/−^ littermate controls.

### RNA extraction and RNAseq analysis

Colonic mucosal tissues were harvested by transecting the colon, washing out contents in ice cold PBS, and scraping mucosa using a glass slide to separate it from the underlying muscle layer. Tissues were harvested and immediately frozen in liquid nitrogen and stored at −80°C. RNA was extracted using TRIzol reagent (Thermo Fisher) as per the manufacturer protocol and stored at −80°C. Preparation of RNA library and transcriptome sequencing was conducted by Novogene Co., LTD (Sacramento, CA).

Analysis of RNAseq was performed according to the methods provided by Novogene as follows (text provided by Novogene): “Raw data (raw reads) of fastq format were firstly processed through in-house perl scripts. In this step, clean data (clean reads) were obtained by removing reads containing adapter, reads containing ploy-N and low-quality reads from raw data. At the same time, Q20, Q30, and GC content the clean data were calculated. All the downstream analyses were based on the clean data with high quality.

Reference genome and gene model annotation files were downloaded from genome website directly. Index of the reference genome was built using Hisat2 v2.0.5 and paired-end clean reads were aligned to the reference genome using Hisat2 v2.0.5. We selected Hisat2 as the mapping tool for that Hisat2 can generate a database of splice junctions based on the gene model annotation file and thus a better mapping result than other non-splice mapping tools. featureCounts v1.5.0-p3 was used to count the reads numbers mapped to each gene. And then FPKM of each gene was calculated based on the length of the gene and reads count mapped to this gene. FPKM, expected number of Fragments Per Kilobase of transcript sequence per Millions base pairs sequenced, considers the effect of sequencing depth and gene length for the reads count at the same time, and is currently the most commonly used method for estimating gene expression levels.

Differential expression analysis of two conditions/groups (two biological replicates per condition) was performed using the DESeq2 R package (1.20.0). DESeq2 provides statistical routines for determining differential expression in digital gene expression data using a model based on the negative binomial distribution. The resulting P-values were adjusted using the Benjamini and Hochberg’s approach for controlling the false discovery rate. Genes with an adjusted P-value ≤0.05 found by DESeq2 were assigned as differentially expressed.”

### Histology scoring

Mouse intestinal tissues were fixed in Methyl Carnoy’s, processed and embedded in paraffin and sectioned at 5μM. Sections were stained by H&E and then scored for histopathology of colitis using methods previously described[10]. Briefly, eight criteria were assessed on a scale of 0–3 to determine the severity of colitis: infiltrate, goblet cell loss, crypt density, crypt hyperplasia, muscle thickening, submucosal infiltrate, and presence of ulceration or abscesses to produce a score ranging from 0 to 24. Each category was assessed with the following criteria: Infiltrate: 0-none, 1-increased presence of inflammatory cells, 2-infiltrates also in submucosa, 3-transmural. Goblet cell loss: 0-none, 1-<10%, 2–10-50%, 3->50%. Crypt density: 0-normal, 1-decreased by <10%, 2-decreased by 10–50%, 3-decreased by >50%. Crypt hyperplasia: 0-none, 1-slight increase in crypt length, 2–2 to 3-fold increase in crypt length, 3-threefold increase in crypt length. Muscle thickening: 0-none, 1-slight, 2-strong, 3-excessive. Submucosal inflammation: 0-none, 1-individual cells, 2-infiltrate, 3-large infiltrates. Crypt Abscess and ulceration was either absent (0) or present (3).

### Immunofluorescence

Methyl Carnoys fixed tissues were processed and sectioned as described above. Following rehydration to PBS, antigen retrieval was performed by boiling in sodium citrate buffer pH6 for 10 minutes followed by 20 minutes cooling, washing in TBS and blocking in TBS + 0.05% Tween-20 (TBST) containing 1% BSA and 5% donkey serum (1hr, RT). Sections were then stained with antibodies to EPCAM and phosphorylated STAT3 in TBST+1% BSA+5%donkey serum (4C, overnight), washed in TBST (3X 5min), incubated in secondary antibodies (1hr, RT), washed in TBST (3 × 5min) and mounted in Prolong Gold media with DAPI. Fluorescent images were acquired on a Leica DM5500 fluorescence microscope.

### Lamina Propria Leukocytes (LPL) Isolation

LPL were isolated from colons as described[11]. Colons along with cecum (without cecal lymphoid patch) were excised, thoroughly cleared of fecal content and residual fat, cut longitudinally, and diced. The sections were washed with several rounds of calcium-magnesium free media with (2%) serum (CMF) to remove feces and mucus. Tissue pieces were shaken at 37°C in CMF with 1mM dithioerythritol (DTE) and 2mM EDTA twice for 20 minutes with agitation after each incubation. Tissue pieces were then rinsed with RPMI and subsequently shaken for ∼1 hour with 100U/ml collagenase (Sigma) and 10U/ml DNase (Sigma) in RPMI with agitation every 20 minutes. Digested tissue sections were then passed through a 70μM filter to eliminate large tissue and collect dissociated cells. Cells were washed in RPMI and centrifuged at 300 xg for 10 minutes. Percoll (40% in RPMI) was then added to cell pellets and centrifuged at 600xg, RT for 20 minutes. Percoll was removed and the cell pellet collected, washed in RPMI, and used for flow cytometry analysis.

### Flow Cytometry

Colon LPLs were suspended in FACS buffer (2%FBS in PBS) and incubated with Fc Block antibody (10min on ice) followed by viability dye eFlour-780 for 30 minutes at 4°C to identify non-viable cells. Cells were then washed with FACS buffer and incubated with antibodies listed in table 1 for 30 minutes at 4°C, washed with FACS buffer and fixed with paraformaldehyde (PFA) (1% PFA in PBS). The fluorescence of individual cells was evaluated through the sample acquisition in a BD LRS-Fortessa X-20 cytometer, analyzed with flow Jo v10.7 and dimensional 2D reduction performed using UMAP[12].

**Table 1:** List of antibodies, fluorophores and reagents used in flow cytometry.

| **Antibody to:** | **Fluorophore** | **Company** | **Identifier** |
| --- | --- | --- | --- |
| CD16/CD32 (FcBlock) | N/A | BD Biosciences | 553142 |
| Viability Dye | eFlour-780 | Thermo Fisher | 65-0865-14 |
| CD45 | BV711 | BioLegend | 103147 |
| CD11b | APC-R700 | BD Biosciences | 564985 |
| CD11c | PE-Cy7 | Thermo Fisher | 25-0114-81 |
| MHCII | PE | BioLegend | 107607 |
| Thy1.2 | FITC | Thermo Fisher | 11-0903-82 |
| SiglecF | eFlour-660 | Thermo Fisher | 50-1702-80 |
| Ly6G | BV650 | BioLegend | 127641 |
| Ly6C | BV421 | BD Biosciences | 562727 |
| NK1.1 | BV785 | Thermo Fisher | 108749 |
| F4/80 | PE-Cy5 | Thermo Fisher | 15-4801-80 |
| CompBeads anti-Rat and Anti-Hamster Igk |  | BD Biosciences | 552845 |
| CompBeads anti-Mouse Igk |  | BD Biosciences | 552843 |

**Table 2:**
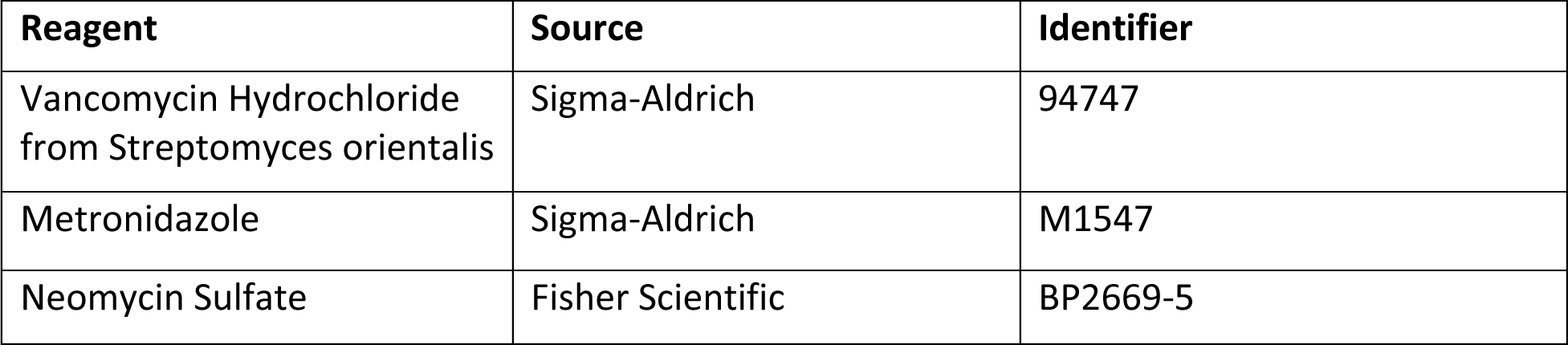
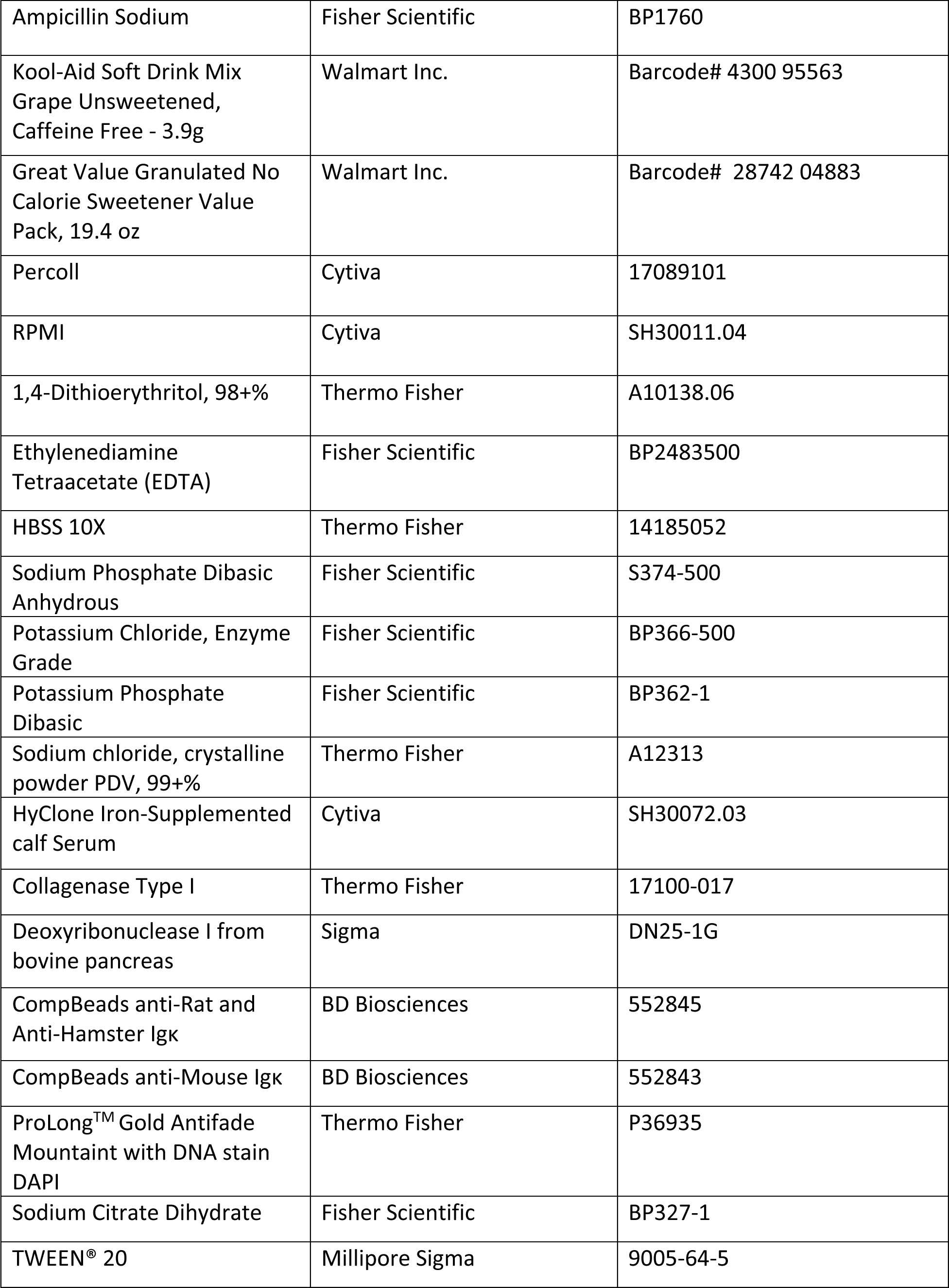
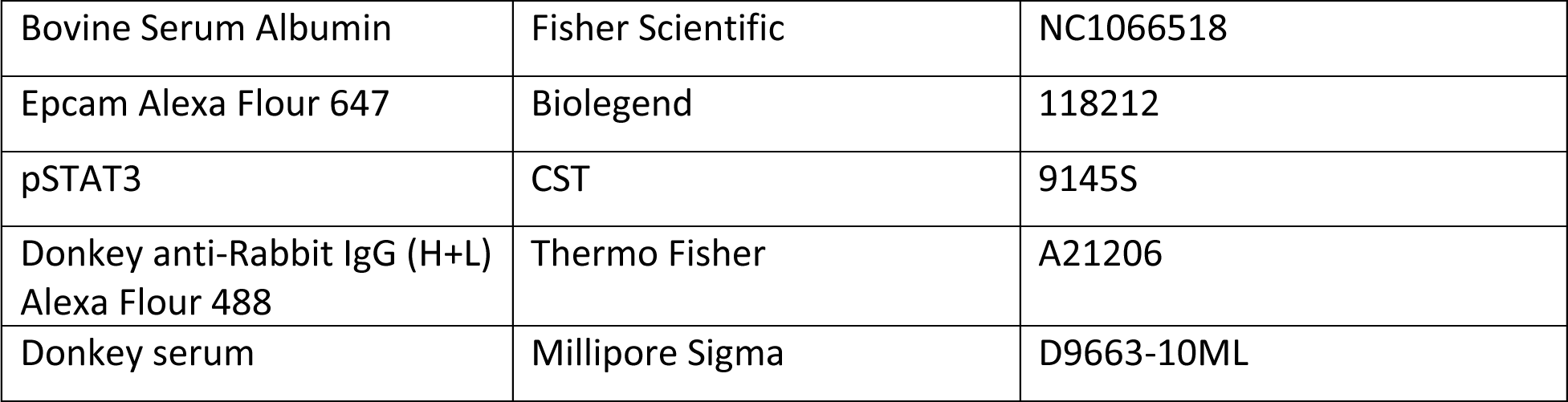
List of reagents.

### Antibiotic treatments

Mice, beginning at 4 weeks of age, were fed a cocktail of four antibiotics, or control water, as described[7]. Ampicillin (0.5g/ml), Neomycin (0.25g/ml), Metranidazole (62.5mg/ml) and Vancomycin (125mg/ml) were provided as a cocktail dissolved in grape KoolAid. Control mice received KoolAid only. Antibiotics were refreshed twice weekly until the mice were 8 weeks of age when colon tissues were collected for histology.

### Statistical analysis

Bar graphs represent the mean plus standard deviation of each group with individual data points representing data from one mouse. Male and female mice were grouped together as there were no significant gender-based differences in the data. Data were analyzed by one way ANOVA with post hoc non-parametric tests for histological scoring (Kruskal-Wallis and Dunn’s multiple comparison) or post-hoc Tukey’s tests for LPL cell counts. Statistical analysis of RNAseq data was performed as described in the Section for RNAseq analysis. Significance was inferred at a p value less than 0.05.

## RESULTS

TRAG mice develop 100% penetrant, early onset colitis in the absence of adaptive antigen receptor expressing cells[6, 7]. We have observed that TRAG mice can develop colitis in the absence of RoRγt, the transcription factor required for ILC3 development[6, 13]. To better characterize and quantify the role of RoRγt in the histopathological features of colitis in TRAG mice, we applied an 8 point histological scoring system[10] comparing RAG1^−/−^, TRAG and Rorc^−/−^ x TRAG histopathology along the proximal to distal axis of the colon. Consistent with prior results, TRAG mice exhibited significantly higher histological scores of colitis compared to RAG1^−/−^ mice, in the cecum, proximal and distal colon (Fig 1). This colitis was also significantly increased in Rorc^−/−^ x TRAG mice, compared to RAG1^−/−^ mice (Fig 1). However, there was a trend toward lower histological scores, which was statistically significant, in the distal colon, of Rorc^−/−^ x TRAG mice, compared to TRAG mice (Fig 1). Thus, the absence of RoRγt ameliorates but does not prevent colitis in TRAG mice.

**Figure 1:**
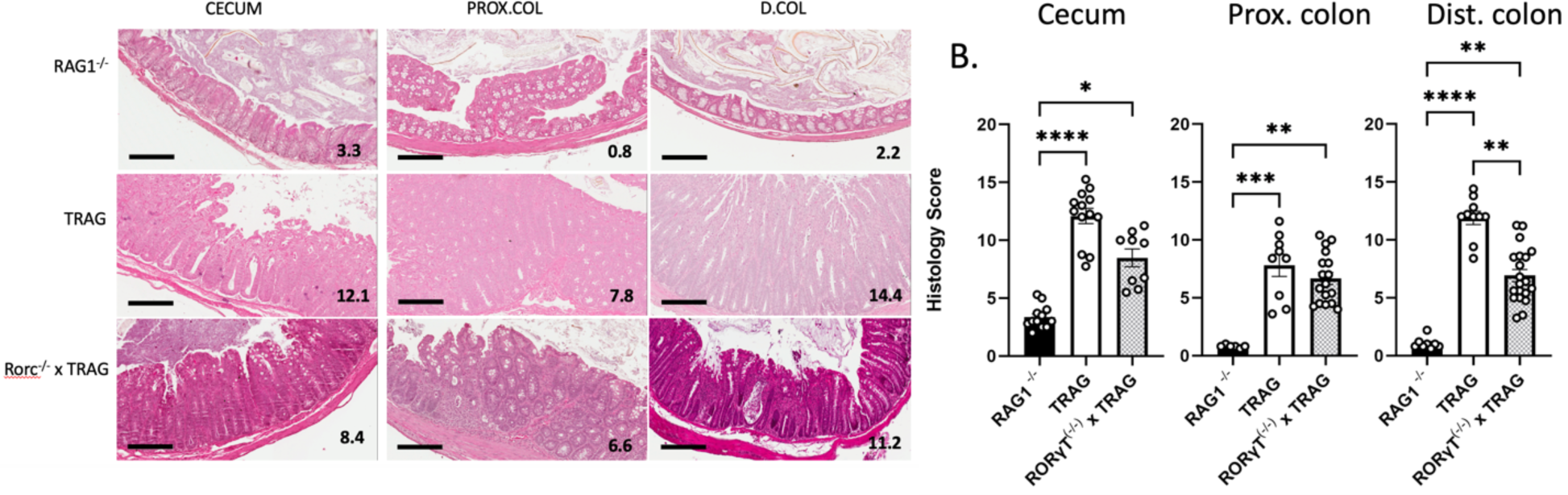
RoRγt is dispensable for colitis in TRAG mice. (**A**) Representative H&E sections of the cecum, proximal colon (PROX.COL) and distal colon (D.COL) of 8-week-old RAG1^−/−^, TRAG and Rorc^−/−^ x TRAG mice. scale bar=200μM. inset numbers are the scores for each section. (**B**) Histological scores of the cecum, proximal colon and distal colon. Each data point represents one mouse. *p<0.05, **p<0.01, ***p<0.005, ****p<0.001.

To assess the cellular innate immune profile of RoRγt-deficient TRAG mice we examined the LPL populations of these mice at 8 weeks of age (Fig 2). Rorc^−/−^ x TRAG mice showed significantly increased numbers of CD45^+ve^ leukocytes in the intestinal mucosa, compared to RAG1^−/−^ mice, and the number of these CD45 leukocytes was comparable between TRAG and Rorc^−/−^ x TRAG mice. Most innate immune cell populations were significantly increased in Rorc^−/−^ x TRAG mice compared to RAG1^−/−^ mice, and were similar in number to those of TRAG mice (Fig 2). One notable exception was that Rorc^−/−^ x TRAG mice had significantly fewer eosinophils than TRAG mice. There were also decreased numbers of neutrophils and ILCs in Rorc^−/−^ x TRAG vs. TRAG mice. This likely reflects the lack of ILC3 in Rorc^−/−^ x TRAG mice and the lack of granulocyte stimulation due the absence of ILC3-derived IL17. Thus, RoRγt^+ve^ cells contribute to the colitis in TRAG mice, especially toward the number of eosinophils in the intestinal mucosa, but RoRγt^+ve^ cells are nonetheless dispensable for colitis in this innate model.

**Figure 2:**
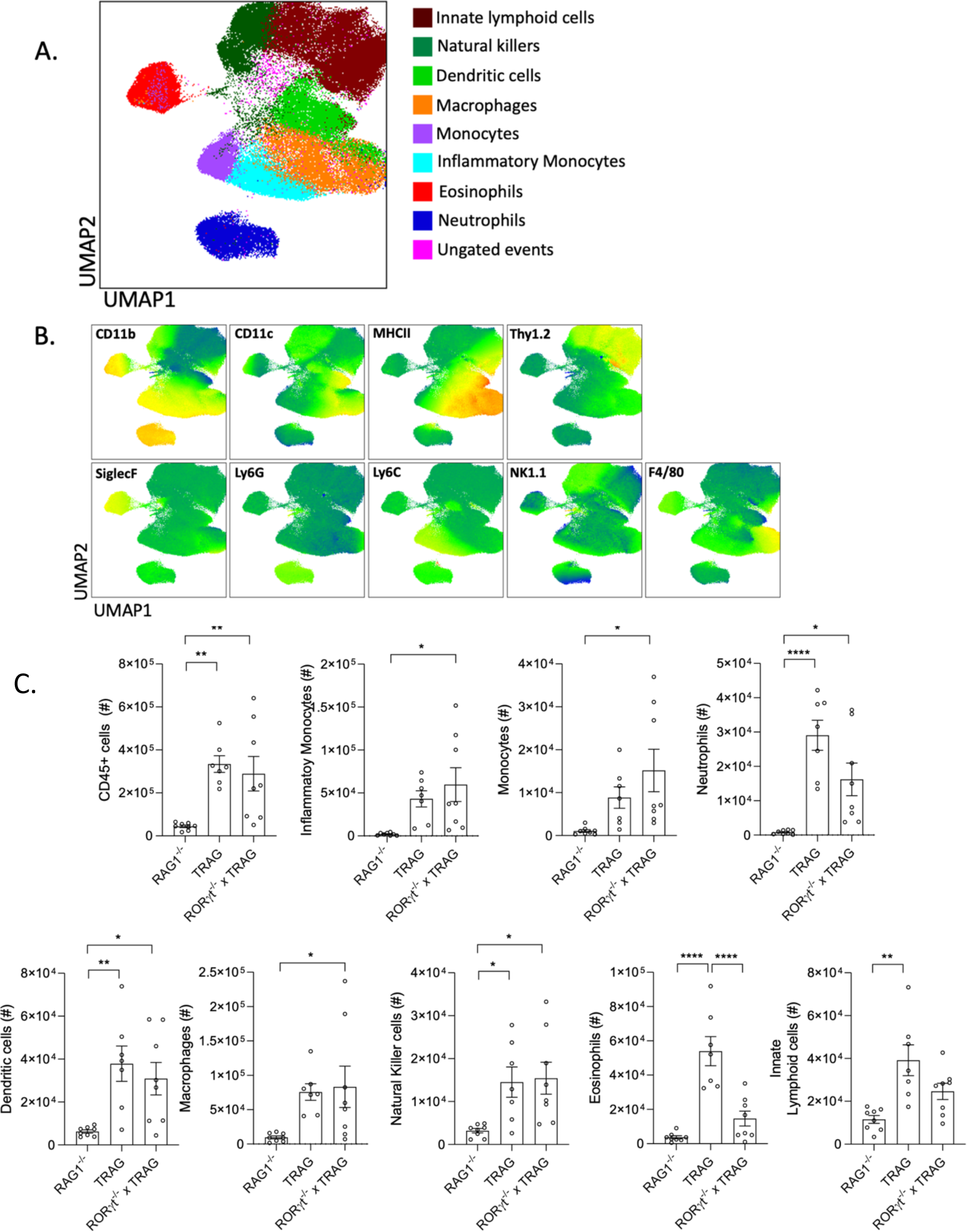
Immune profile of TRAG mice lacking RoRγt. (A,B) UMAP plots with markers of immune cell populations isolated from the lamina propria of 8-week-old RAG1^−/−^, TRAG and Rorc^−/−^ x TRAG mice. (C) Cell counts from LPL preparations of 8-week-old RAG1^−/−^, TRAG and Rorc^−/−^ x TRAG mice. Each data point represents one mouse. *p<0.05, **p<0.01, ***p<0.005, ****p<0.001.

To understand the molecular features of colitis in Rorc^−/−^ x TRAG mice, we performed RNAseq of TRAG and Rorc^−/−^ x TRAG colonic mucosal scrapings. TRAG and Rorc^−/−^ x TRAG colons had 451 and 586 uniquely expressed genes, respectively, and 12064 genes were expressed in both gentotypes (Fig 3). There were 1876 differentially expressed genes, with 972 increased in expression in TRAG vs. Rorc^−/−^ x TRAG mice, and 904 genes exhibiting decreased expression. GO enrichment analysis indicated that biological processes were enriched for neutrophil and leukocyte chemotaxis and regulation of inflammation in TRAG vs. Rorc^−/−^ x TRAG mice (Fig 3). Molecular functions enriched in TRAG vs. Rorc^−/−^ x TRAG mice included cytokine and chemokine signaling. KEGG enrichment analysis of TRAG vs. Rorc^−/−^ x TRAG mucosa found that cytokine-receptor, IL17 signaling, and inflammatory bowel disease were the most significantly enriched pathways (Fig 3). Other enriched pathways included TNF signaling and NFkappaB signaling. Similarly, the reactome enrichment analysis comparing TRAG vs. Rorc^−/−^ x TRAG implicated cytokine signaling and neutrophil degranulation but also antimicrobial peptides and IL-6 family signaling (Fig 3). Together these transcriptome results are consistent with reduced colitis observed in the histopathology of Rorc^−/−^ x TRAG mice, compared to TRAG mice, and especially to the reduced granulocyte populations in Rorc^−/−^ x TRAG intestinal mucosa.

**Figure 3:**
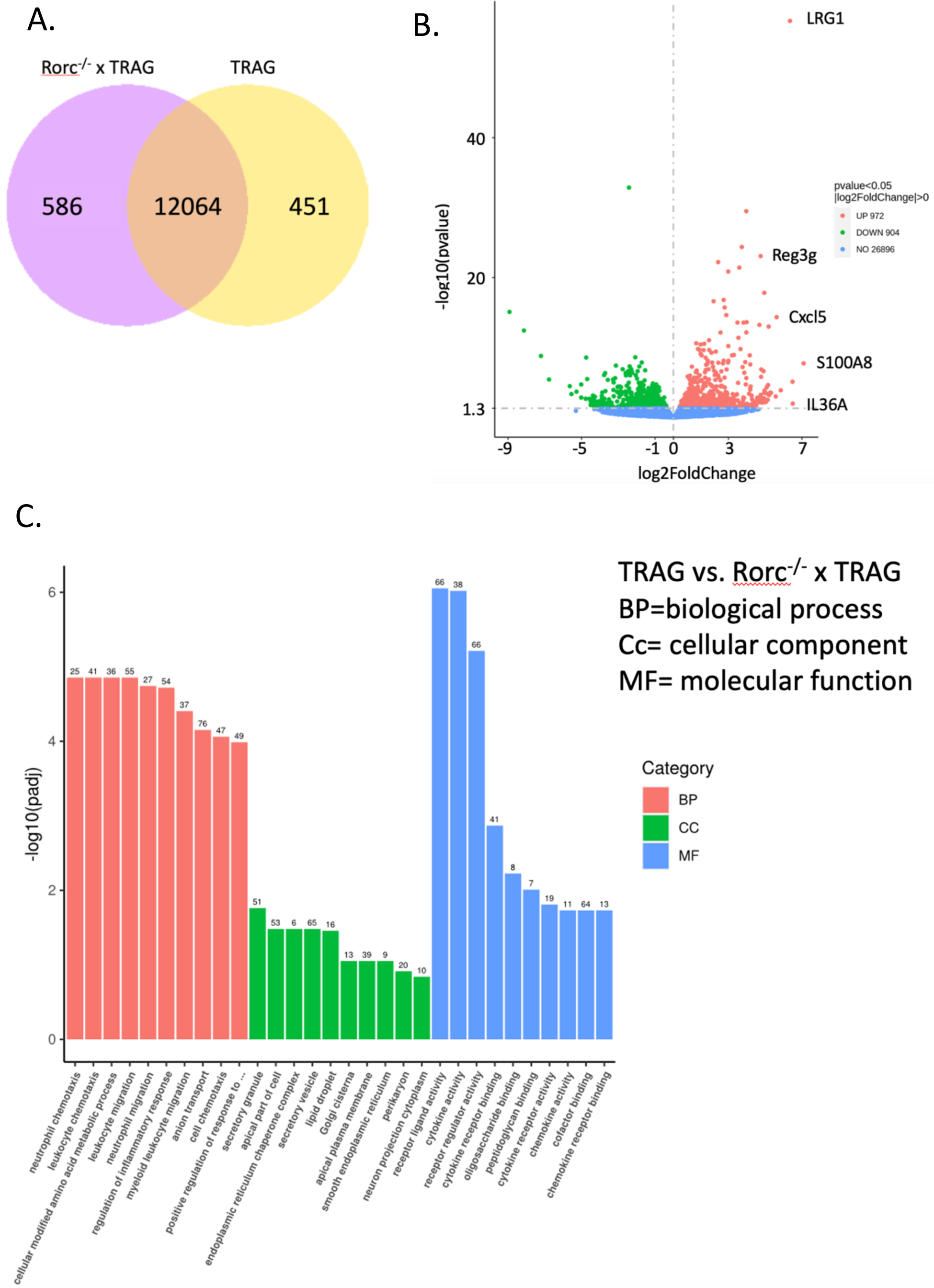
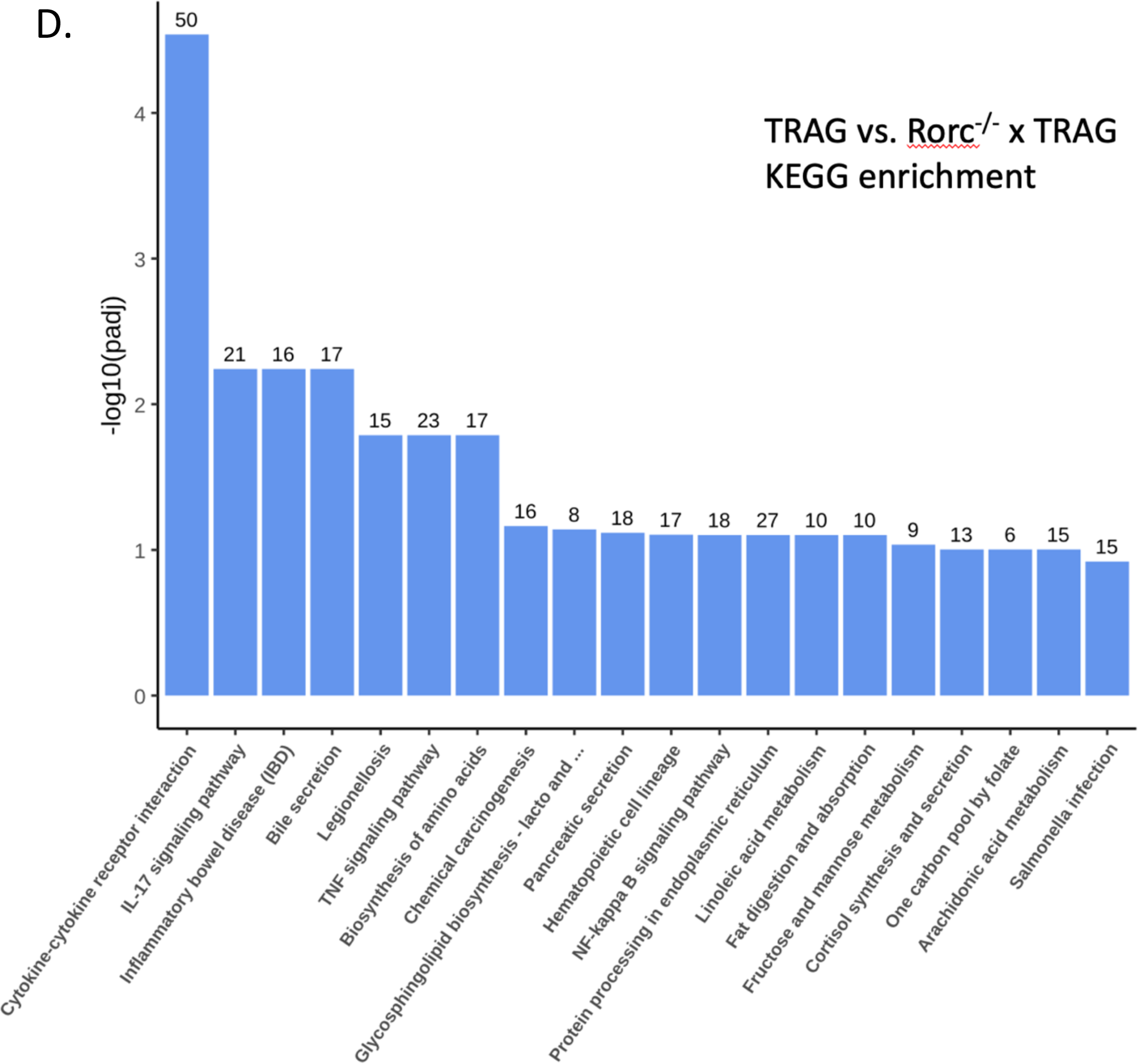
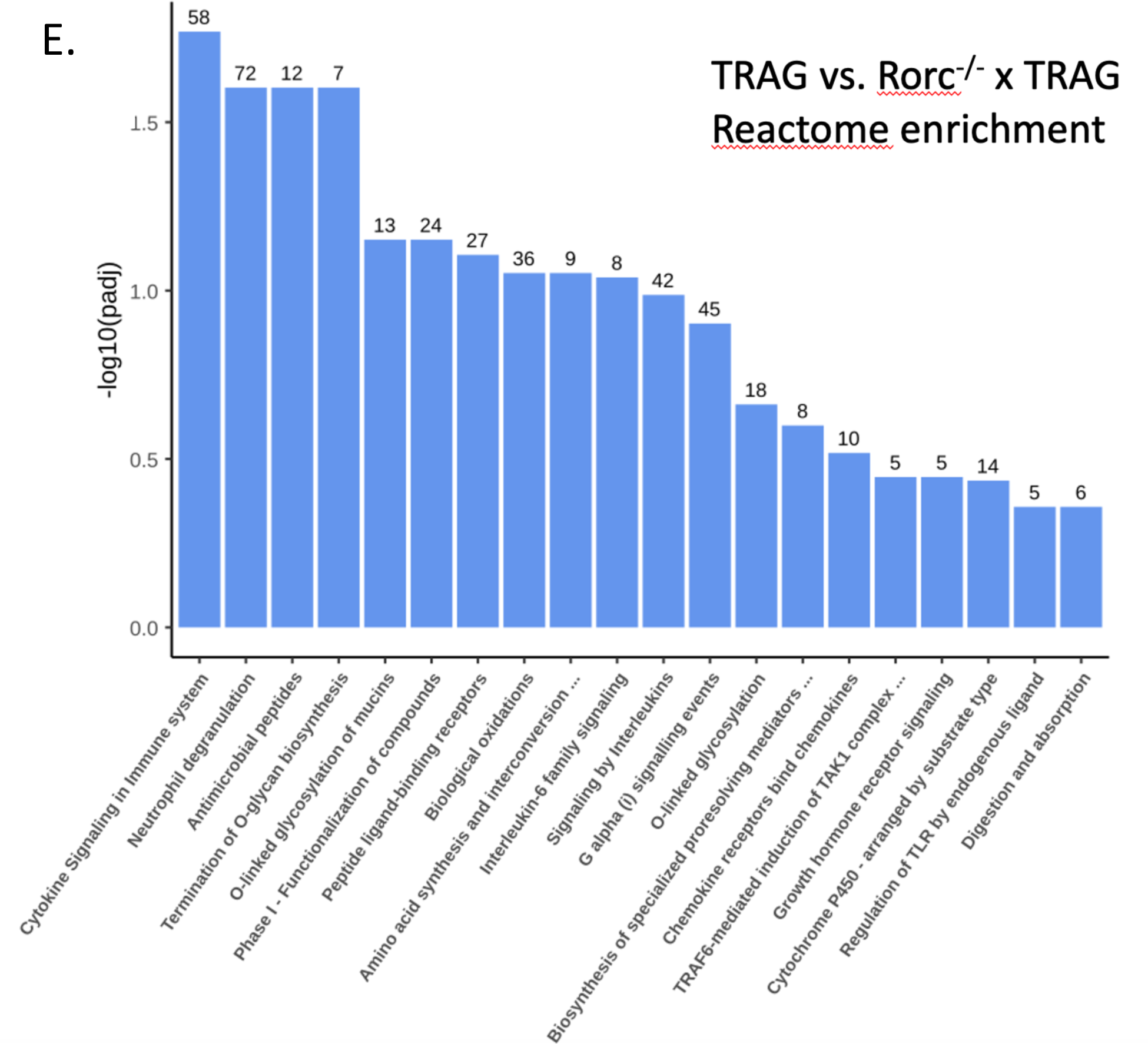
RNA profiles of TRAG and Rorc^−/−^ x TRAG colonic mucosa. **(A)** Venn diagram of shared and unique transcripts. **(B)** Volcano plot of RNA with increased and decreased expression in TRAG vs. Rorc^−/−^ x TRAG colonic mucosa. **(C)** GO **(D)** KEGG and **(E)** Reactome enrichment analysis of TRAG vs. Rorc^−/−^ x TRAG colon mucosa.

RoRγt expressing ILC3 produce IL17 and IL22, both of which can contribute to eosinophilia and granulocyte recruitment, which may explain the reduced the reduced numbers of granulocytes in the mucosa of Rorc^−/−^ x TRAG mice compared to TRAG mice[14, 15]. In addition, expression of other eosinophil activating cytokines like IL33 and IL36a [16–19] were reduced in Rorc^−/−^ x TRAG colons (Fig 4). Lastly, expression of chemokines that are produced by activated eosinophils, including cxcl1 and cxcl5 [16, 20] were reduced in Rorc^−/−^ x TRAG colons, consistent with the significant reduction of eosinophil numbers in the mucosa of Rorc^−/−^ x TRAG mice (Figs 2, 4). The reduced expression of cytokines that induce eosinophilia and the reduced expression of cytokines made by activated eosinophils is consistent with the significant reduction of eosinophils we observed in the colons of Rorc^−/−^ x TRAG mice compared to TRAG mice. Despite this significant reduction in eosinophils, Rorc^−/−^ x TRAG mice do not show improved immune profiles of other inflammatory cells or amelioration of their colitis. Thus, eosinophils numbers are reduced but this does not ameliorate colitis in TRAG mice lacking RoRγt.

**Figure 4:**
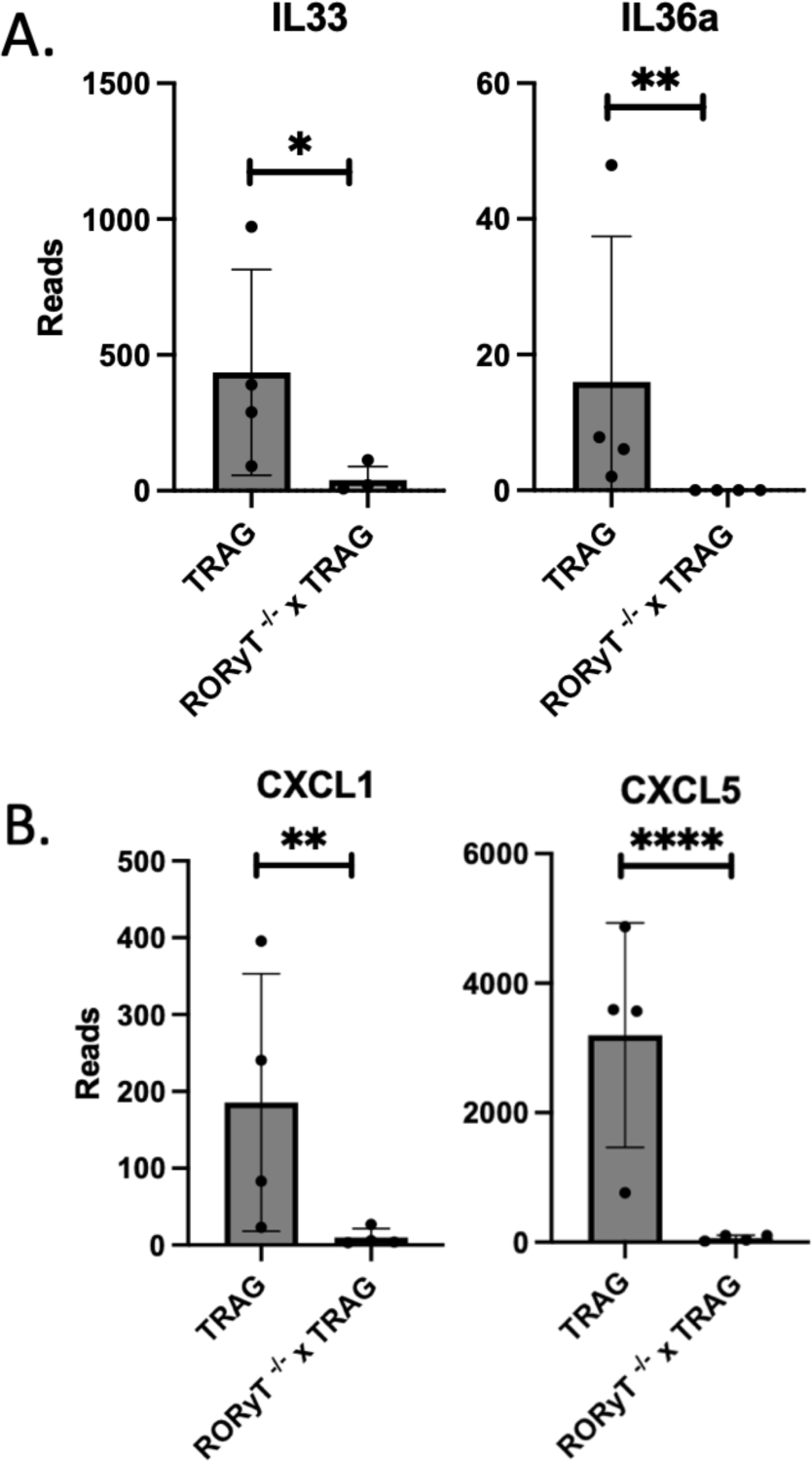
Reduced expression of cytokines that (A) induce eosinophilia and (B) are made by eosinophils in the colon mucosa of Rorc^−/−^ x TRAG vs. TRAG mice. *p<0.05, **p<0.01, ***p<0.001.

ILC3 derived IL17 and IL22 are also protective cytokines that promote antimicrobial gene expression, the latter through IL22R-induced STAT3 signaling[21–24]. Consistent with this, reactome enrichment analysis of TRAG vs. Rorc^−/−^ x TRAG mice identified antimicrobial peptides and IL-6 family signaling, which are STAT3 mediated pathways (Fig 3). IL22 produced by ILC3 drives STAT3 dependent antimicrobial gene expression that persists in RAG1^−/−^ small intestines after colonization with microbes[21]. We therefore examined STAT3 activation in TRAG vs. Rorc^−/−^ x TRAG colons. Consistent with observations in RAG1^−/−^ small intestine[21], intestinal epithelial cells of 8-week-old TRAG colons exhibited positive nuclear localization of phospho-STAT3, indicating that STAT3 was induced in these cells (Fig 5). Rorc^−/−^ x TRAG mice did not exhibit this pattern of phospho-STAT3 nuclear localization. These observations are consistent with the transcriptomic profile indicating reduced IL6 family cytokine signaling in Rorc^−/−^ x TRAG vs. TRAG mice, and consistent with prior reports indicating that ILC3-derived IL22 induces STAT3 in immunodeficient mice[21].

**Figure 5:**
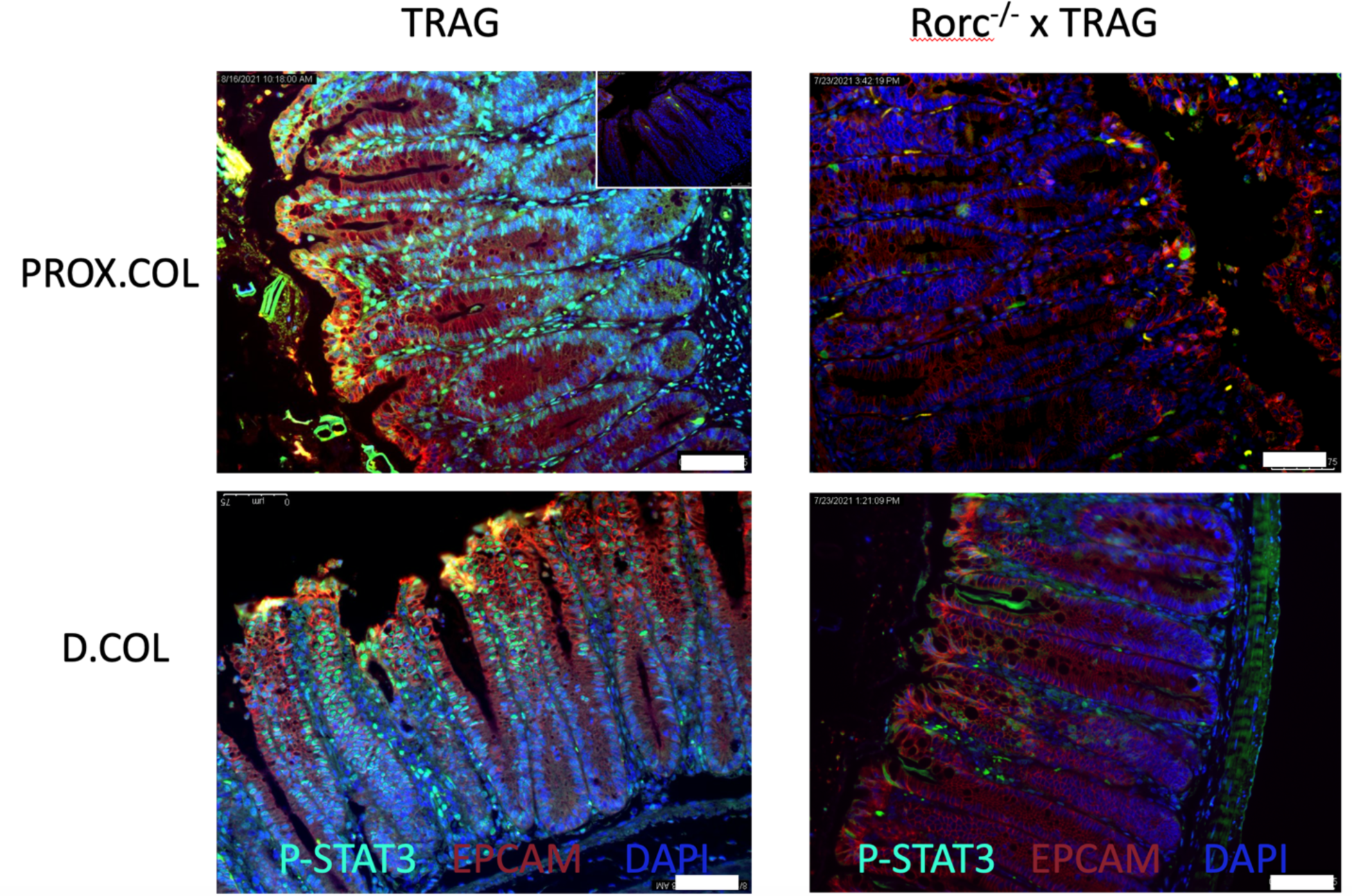
Reduced activation of STAT3 in TRAG mice lacking RoRγt. Immunolocalization of phospho-STAT3 in nuclei of epithelial cells from the distal and proximal colon of TRAG mice vs. Rorc^−/−^ x TRAG mice. (inset, top right=no primary control). Scale bar = 75μM.

ILC3 derived IL22 induces phosho-STAT3 in intestinal epithelial cells resulting in transcription of antimicrobial genes[21]. We therefore examined the antimicrobial gene and STAT3 dependent transcripts in Rorc^−/−^ x TRAG vs. TRAG colonic mucosa. Rorc^−/−^ x TRAG colon mucosa expressed significantly less Reg3β, Reg3γ, S100A8 and S100A9, and less IL22 than TRAG colon mucosa (Fig 6). Other STAT3 induced genes known to be induced by IL22 in intestinal epithelial cells were also markedly reduced in Rorc^−/−^ x TRAG colons, including Muc1, Bcl2l15, OSMR, LRG1, Fut2, Duox2, Sprr2h, NOS2, and SOCS3 (Fig 6). Intestinal epithelial genes involved in mucosal repair or epithelial homeostasis that displayed increased expression in Rorc^−/−^ x TRAG mucosa vs. TRAG mucosa included SELENOP, Sult1a1, Guca2a, SSTR1, FABP6, and TSKU. Thus, the absence of RoRγt leads to reduced expression of STAT3 dependent genes and compensatory gene signature consistent with mucosal repair.

**Figure 6:**
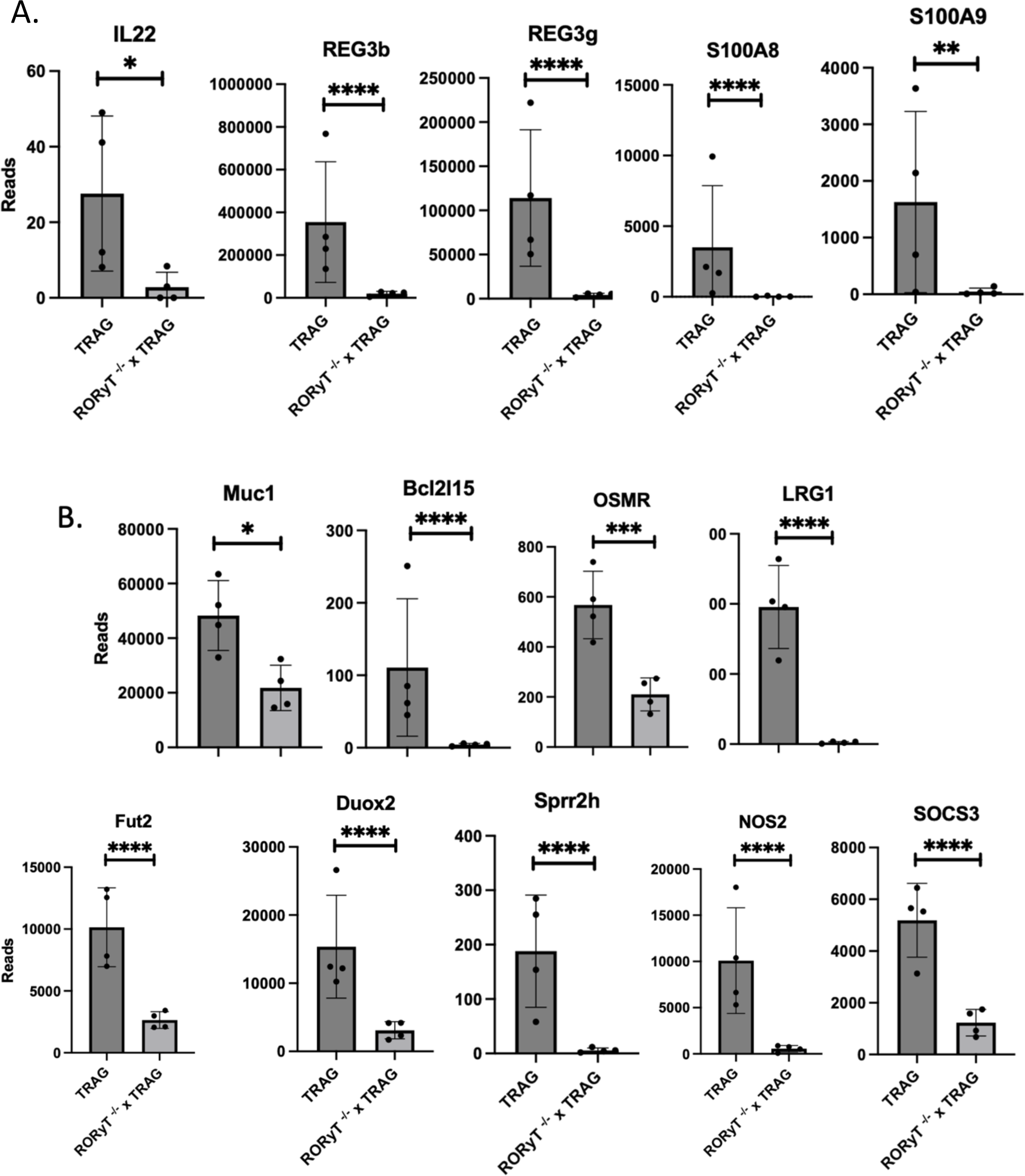
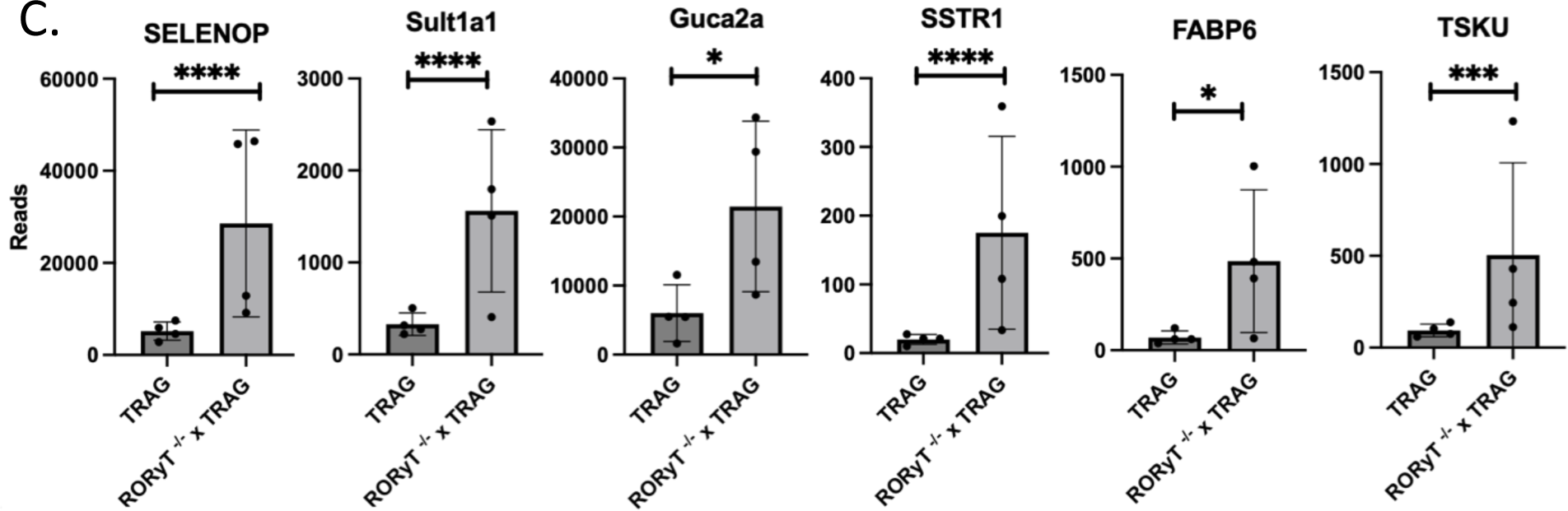
Gene expression changes in Rorc^−/−^ x TRAG vs TRAG colon mucosa. Decreased expression of (A) IL22 and antimicrobial factors, and (B) decreased expression of other IL22-induced genes. (C) Increased expression of mucosal repair related genes in the mucosa of Rorc^−/−^ x TRAG colon. Each data point represents one mouse. *p<0.05, **p<0.01, ***p<0.005, ****p<0.001

The reduced expression of antimicrobial genes in Rorc^−/−^ x TRAG intestinal mucosa suggested that one reason Rorc^−/−^ x TRAG mice develop colitis may be due to microbes populating the gut. To test this, we treated 4-week-old Rorc^−/−^ x TRAG mice with antibiotics and assessed colitis at 8 weeks of age. Colitis was significantly attenuated by antibiotic treatment in the cecum, proximal colon, and distal colon of Rorc^−/−^ x TRAG mice (Fig 7). Thus, colitis in Rorc^−/−^ x TRAG mice is driven by microbes and can be prevented by administration of antibiotics. Together, these results reflect the dual nature of ILC3 in the intestine, promoting colitis by activating innate inflammation but preventing colitis by inducing antimicrobial responses through the IL22-STAT3 pathway in intestinal epithelial cells.

**Figure 7:**
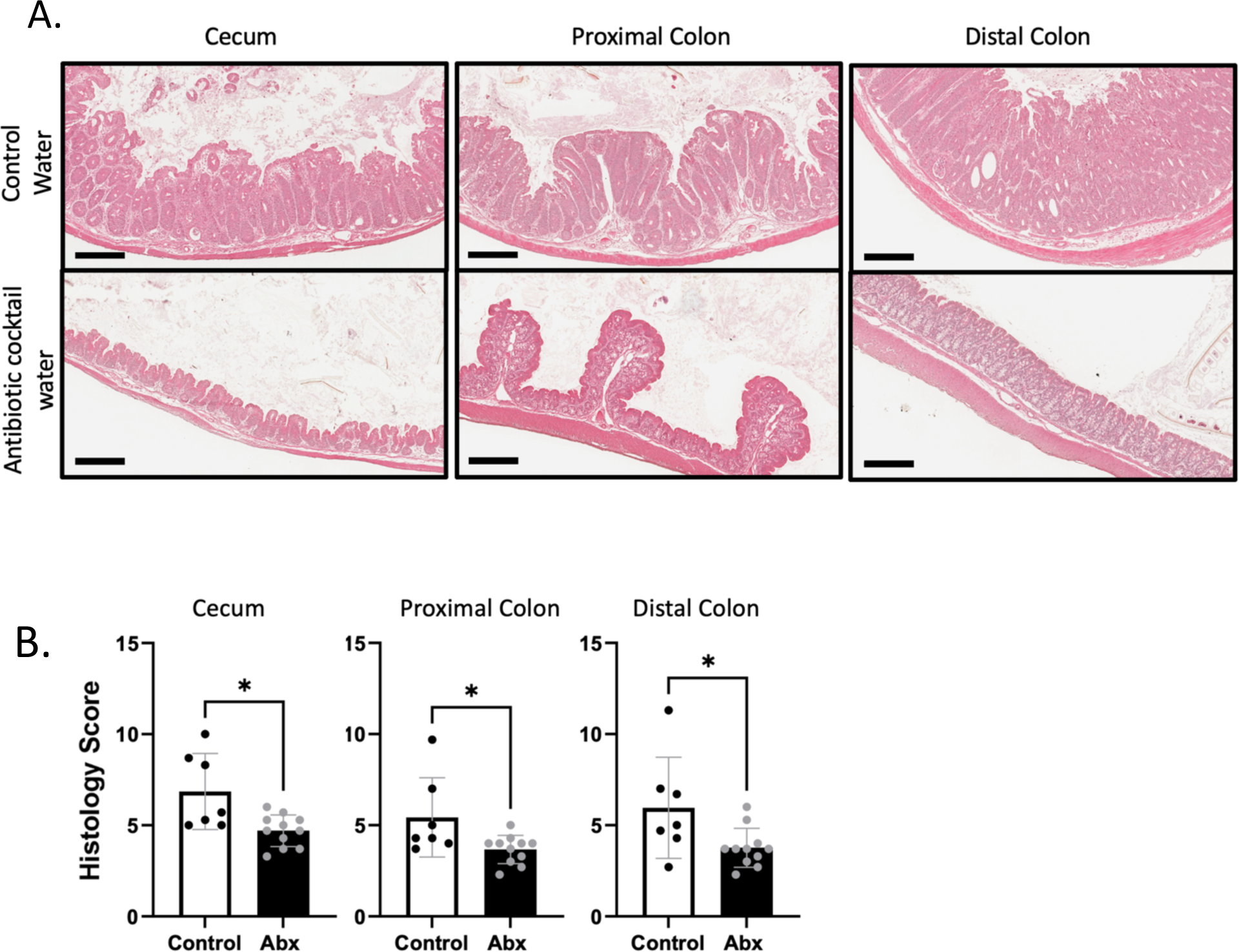
Amelioration of colitis in Rorc^−/−^ x TRAG mice treated with antibiotics. (A) Histological images of cecum, proximal colon and distal colon of 8-week-old Rorc^−/−^ x TRAG mice treated with control (grape KoolAid; upper panels) water or antibiotic cocktail water (lower panels) for 4 weeks. (B) Histological scores of tissues from Rorc^−/−^ x TRAG mice treated with control water or antibiotic cocktail (Abx). *p<0.05

## DISCUSSION

Our results indicate that RoRγt contributes to inflammation and innate immune colitis in TRAG mice primarily in the distal colon. RoRγt is especially important for elevated numbers of eosinophils and, to a lesser extent neutrophils and ILCs in the intestinal mucosa of TRAG mice. Eosinophils, and not neutrophils, driven by the IL23-GMCSF axis have been implicated in colitis in both the T cell transfer and the H. hepaticus + anti-IL10R models of IBD[25]. The eosinophil driven colitis in those models exhibited increased expression of mucosal GMCSF, eotaxin, and RANTES but not IL5[25]. We did not observe changes in expression of any of these cytokines between TRAG and Rorc^−/−^ x TRAG mice, despite significant decreases in eosinophils in the latter. This may be due to the fact that IL5 and GMCSF are predominantly T cell derived cytokines and the TRAG model lacks all adaptive immunity. However, ILC3 cells, which are absent in Rorc^−/−^ x TRAG mice, are critical for the production of GMCSF driven innate colitis in the anti-CD40 or H hepaticus infection of RAG^−/−^, which are innate models of IBD[26]. It is possible that a lack of ILC3 derived GMCSF accounts for the reduced eosinophilia we observed in Rorc^−/−^ x TRAG colon mucosa. There was also a significant decrease in expression of IL33, IL36a and IL36g in Rorc^−/−^ x TRAG mucosa, and these cytokines can induce eosinophil accumulation/activation[16, 17, 19], but these cytokines are not considered RoRγt or ILC3 dependent. Another explanation for reduced eosinophils in the mucosa of Rorc^−/−^ x TRAG mice may be that RoRγt plays a direct role in the function of eosinophils. Expression of RoRγt leading to IL17 production by eosinophils has been reported for eosinophils that accumulate in the lungs in models of acute and allergic aspergillosis[27]. Whether RoRγt plays a similar role in gut eosinophils remains to be determined. Our data indicate that eosinophil accumulation, but not colitis, are dependent on RoRγt in TRAG mice suggesting that eosinophil-independent and RoRγt-independent mechanisms drive the colitis in TRAG mice.

RoRγt is critical for the development of ILC3, which have been implicated as drivers of innate models of IBD[5, 28]. Despite this, RoRγt is largely dispensable for innate colitis in TRAG mice. This may represent the dual nature of ILC3 having both proinflammatory and antimicrobial roles in the intestine[29]. ILC3 produce IL22 which is a key cytokine for activation of epithelial cell STAT3 and induction of antimicrobial responses[21]. Indeed, Rorc^−/−^ x TRAG mice displayed significant reductions in pSTAT3 in the intestinal epithelium along with reduced expression of multiple antimicrobial factors including Reg3γ, Reg3γ and calprotectin (S100A8 and S100A9). The reduction of these antimicrobial defenses may contribute to the colitis in Rorc^−/−^ x TRAG mice, as antibiotic treatment was able to reduce colitis in these mice. This microbial driven inflammation may also be exacerbated by expression of TNFAIP3 in IEC which is permissive for invasion of the inner mucus layer of the colon by Actinobacteria and Gammaproteobacteria[7, 8].

In addition to the development of ILC3, RoRγt regulates the expression of IL23R and thus responsiveness of intestinal ILC3 to IL23[28]. In the anti-CD40 antibody and *H. hepaticus* induced models of innate colitis, RoRγt-dependent ILC3 are required for development of colitis[5]. Although we observed some attenuation of colitis in the distal colon of Rorc^−/−^ x TRAG mice, the colitis was more severe compared to RAG1^−/−^ distal colons. In addition, there was no significant reduction in colitis severity in the cecum or proximal colon of Rorc^−/−^ x TRAG mice compared to TRAG mice. The innate colitis in anti-CD40 or *H. hepaticus* treated mice is acute and systemic, whereas the colitis in TRAG mice is chronic and limited to the colon. The systemic inflammation in these other models of innate colitis is driven by IL12, whereas the mucosal inflammation is driven by IL23[30]. The persistence of colitis in Rorc^−/−^ x TRAG mice in the absence of systemic inflammation and the absence of ILC3 suggests that colitis in TRAG mice is driven by cytokines other than IL12 or IL23. The cellular mechanism of *H. hepaticus* induced innate colitis is not known, but anti-CD40 induces innate colitis by activating CD40 expressing antigen presenting cells (primarily dendritic cells) and can occur in germ-free mice[30]. Whether TRAG or Rorc^−/−^ x TRAG mice develop colitis in germ free conditions is not known, but the colitis in TRAG or Rorc^−/−^ x TRAG mice is prevented by antibiotics suggesting that microbes are required for colitis in this model[6]. Thus, the chronic, spontaneous, innate colitis of TRAG mice is distinct from the colitis observed in other innate models and this is also reflected in the RoRγt-independent colitis of this model.

Although the effects of RoRγt deletion are likely due to the loss of ILC3 in Rorc^−/−^ x TRAG mice, it should be noted that RoRγt plays roles in other cell types. A subset of neutrophils expresses RoRγt and rapidly produces IL17 in response to IL6 and IL23[31]. These neutrophils are not dependent on RoRγt for development but play roles in responses to fungal pathogens and in inflammatory diseases. Whether RoRγt^+ve^ neutrophils play a role in IBD is not known. Additionally, RoRγt^+ve^ AIRE^+ve^ antigen presenting cells distinct from ILC3 have been described[28, 32]. These cells promote T cell responses to *Candida* but they are not typically found in the intestine[28]. Since the Rorc^−/−^ x TRAG model has no antigen receptor bearing cells, it is unlikely that these RoRγt^+ve^ APCs contribute to the innate inflammation in this model of colitis.

We observed a gene signature in Rorc^−/−^ x TRAG intestine consistent with epithelial cell injury and repair. There was increased expression in Rorc^−/−^ x TRAG mucosa, compared to TRAG mucosa, of SELENOP, TSKU, Sult1a1 and Guca2a, which are signatures of wound healing or intestinal epithelial responses to damage and are potential biomarkers of CD and UC[33–37]. Both IL17 and IL22 promote epithelial cell integrity and repair and the loss of RoRγt^+ve^ ILC3, a major source of IL17 and IL22 in the gut, likely leads to increased intestinal damage and the compensatory induction of these would response pathways.

RoRγt, or genes regulated by RoRγt, are promising targets for therapy but results from this study suggest that other RoRγt-independent mechanisms can contribute to innate immune mediated colitis. Thus, future efforts to identify RoRγt -independent mechanisms that contribute to innate immune mediated colitis will provide important additional information to help guide therapies to control innate immune inflammation in IBD.

